# Cdc24, the source of active Cdc42, transiently interacts with septins to create a positive feedback between septin assembly and bud site formation in yeast cells

**DOI:** 10.1101/719815

**Authors:** Julian Chollet, Alexander Dünkler, Anne Bäuerle, Laura Vivero-Pol, Thomas Gronemeyer, Nils Johnsson

**Author notes:** equal contribution (Julian Chollet, Alexander Dünkler). Corresponding author, Contact: Nils Johnsson, Institute of Molecular Genetics and Cell Biology, Department of Biology, Ulm University, James-Franck-Ring N27, D-89081 Ulm, Germany.

## Abstract

Yeast cells select at the beginning of each cell cycle the position of their new bud. The recruitment of the septins to this prospective bud site (PBS) is one of the critical events in a complex assembly pathway that culminates in the outgrowth of a new daughter cell. The septin-rods follow hereby the high concentration of Cdc42_GTP_ that is generated by the focused location of its GEF Cdc24. We show that Cdc24 not only activates Cdc42 but temporarily interacts shortly before budding with Cdc11, the subunit that caps septin rods at its both ends. Mutations in Cdc24 that reduce the affinity to Cdc11 impair septin assembly and decrease the stability of the polarity patch. The interaction between septins and Cdc24 thus reinforces bud assembly at sites where septin structures are formed. Once the septins polymerize into the ring, Cdc24 transfers to its center and directs from there the further outgrowth of the membrane.

## Introduction

Yeast cells grow and divide by forming a bud at the beginning of each cycle. The bud matures through polarized growth into the future daughter cell. The formation of the new bud requires in a first step a focused assembly of a complex set of proteins and lipids to a preselected point of the plasma membrane (Howell and Lew, 2012; Howell et al., 2009). The position of this prospective bud site (PBS) is selected by landmark proteins that locally activate the Ras-like Rsr1 (Park et al., 1997; Bender and Pringle, 1989; Chant and Herskowitz, 1991; Kang et al., 2001). Rsr1_GTP_ then recruits and possibly stimulates Cdc24, the GEF for the Rho GTPase Cdc42 (Park et al., 1997; Shimada et al., 2004). Cdc42_GTP_ directly binds and activates approximately a dozen of effector proteins whose combined activities push the plasma membrane and the cell wall to initiate bud formation and growth (Bose et al., 2001; Evangelista et al., 1997; Irazoqui et al., 2003; Chiou et al., 2017). Many more proteins are recruited in addition as direct or indirect consequences of Cdc42 activation (Pruyne et al., 2004). In spite of this massive protein and membrane influx PBS assembly stays remarkably focused. Only one bud with a base of less than 2 µm in diameter is formed. The association of the landmark proteins with the previous cell division site ensures that the new bud of haploid cells forms always next to it (Chant and Herskowitz, 1991).

A critical step in the PBS assembly involves the recruitment of the septins. The four septin-units Cdc11, Cdc12, Cdc3 and Cdc10 form octameric rods that polymerize at the bud neck into filaments and higher-order structures like rings and collars (Bertin et al., 2008; Marquardt et al., 2019). Cdc11 is the subunit that caps the rod at its both ends and thus plays a critical role in filament formation (Garcia et al., 2011; Brausemann et al., 2016). During mitosis the septins split into two rings and enclose the space where cytokinesis and abscission occurs (Marquardt et al., 2019). After abscission septins are disassembled from the site of cytokinesis and transferred in a Cdc42_GTP_-dependent manner to the new PBS (Gladfelter et al., 2002; Caviston et al., 2003). The probably not fully structured patch of octameric septin rods then polymerizes again into the ring-forming filaments. The ring remains at the base of the bud during further membrane and cell wall growth and restricts the free movement of the plasma membrane between mother and daughter cell (Barral et al., 2000). The mechanisms of septin recruitment and assembly are not fully understood (Marquardt et al., 2019). A central role is played by the PAK Cla4 and the paralogous proteins Gic1 and Gic2 (Kadota et al., 2004). Gic1 and Gic2 are effectors of Cdc42_GTP_ and are discussed to facilitate in their GTP-bound state the transfer and the incorporation of septin rods at the PBS (Iwase et al., 2006; Okada et al., 2013; Sadian et al., 2013). Cla4 phosphorylates certain subunits of the septins and like the PAK Ste20 is thought to be stimulated by its direct interaction with Cdc42_GTP_ (Versele and Thorner, 2004; Lamson et al., 2002). A loss of the kinase impairs septin localization and its assembly into rings (Weiss et al., 2000; Kadota et al., 2004). These effects are aggravated by the simultaneous deletion of members of the polarisome, a complex of proteins that catalyze the formation of actin filaments at sites of polar growth (Kadota et al., 2004).

The assembly of the PBS is under cell cycle control and requires activation of CDK in late G1 (Witte et al., 2017; Lai et al., 2018; McCusker et al., 2007; Atkins et al., 2013; Gulli et al., 2000; Moran et al., 2019). CDK also influences the order of assembly as the time interval between Bem1-Cdc24 assembly and arrival of the septins can be shortened by pre-activation of CDK (Lai et al., 2018). The enforcement of a linear order of biochemical events might be necessitated by the complexity of PBS assembly. Assembly pathways can equip structure formation with robustness and flexibility through positive or negative inputs from previous or simultaneously occurring steps. As the only source of active Cdc42, the localization and regulation of Cdc24 is especially critical for bud site assembly (Sloat and Pringle, 1978; Sloat et al., 1981). Consequently, important positive and negative feedback loops convene on Cdc24’s GEF activity (Kozubowski et al., 2008; Kuo et al., 2014; Goryachev and Pokhilko, 2008; Wedlich-Soldner et al., 2004; Wai et al., 2009; Witte et al., 2017; Smith et al., 2013; Shimada et al., 2004; Butty et al., 2002). Cdc24 is constitutively bound to the scaffold Bem1 through their C-terminally located PB domains (Ito et al., 2001; Peterson et al., 1994). Bem1 not only determines the localization of Cdc24 but also influences its activity either directly or indirectly through its bound partner proteins (Witte et al., 2017; Woods et al., 2015; Smith et al., 2013; Rapali et al., 2017).

This work characterizes a novel interaction between the septins and the Cdc24-Bem1 complex. The interaction is part of a feedback that ties the continuous production of active Cdc42_GTP_ to successful septin assembly.

## Results

### The Cdc24-Bem1 complex binds the septin subunit Cdc11

A more complete understanding of its interactome might reveal how Cdc24 is regulated and how its localization is defined during the cell cycle. A library-based Split-Ubiquitin screen identified the septin subunit Cdc11 as novel interaction partner of Cdc24 (Johnsson and Varshavsky, 1994). We confirmed our initial finding by testing the interactions of Cdc24 extended by the C_ub_-RUra3 module (Cdc24CRU) against an array of 548 yeast strains that expresses besides N_ub_-Cdc11 N_ub_ fusions to proteins known to play roles in polarity establishment and polar growth, stress response, cytokinesis and further cellular processes (Hruby et al., 2011; Wittke et al., 1999). The array identified known and novel binding partners of Cdc24 (Fig. 1A, Table 1, Fig. S1), and proofed the specificity of the Cdc24-Cdc11 interaction as none of the other four N_ub_-labeled members of the mitotic septins generated an interaction signal with Cdc24CRU (Fig. 1B). Cdc24 forms a constitutive complex with Bem1. To decide whether Cdc11 interacts directly with Cdc24 or indirectly through Bem1, we tested N_ub_-Cdc11 against Cdc24CRU in the presence and absence of Bem1. The deletion of *BEM1* is lethal in the strain JD47, but *Δbem1-*cells can be kept viable though the simultaneous deletion of the Cdc42 GAP *BEM3* (*Δbem1 Δbem3)* (Laan et al., 2015; Dowell et al., 2010). The test clearly demonstrates that Cdc11, in contrast to Boi1 and Boi2, belongs to those proteins (Rga2, Rsr1) that do not require Bem1 for their interaction with Cdc24 (Fig. 1C).

**Figure 1.**
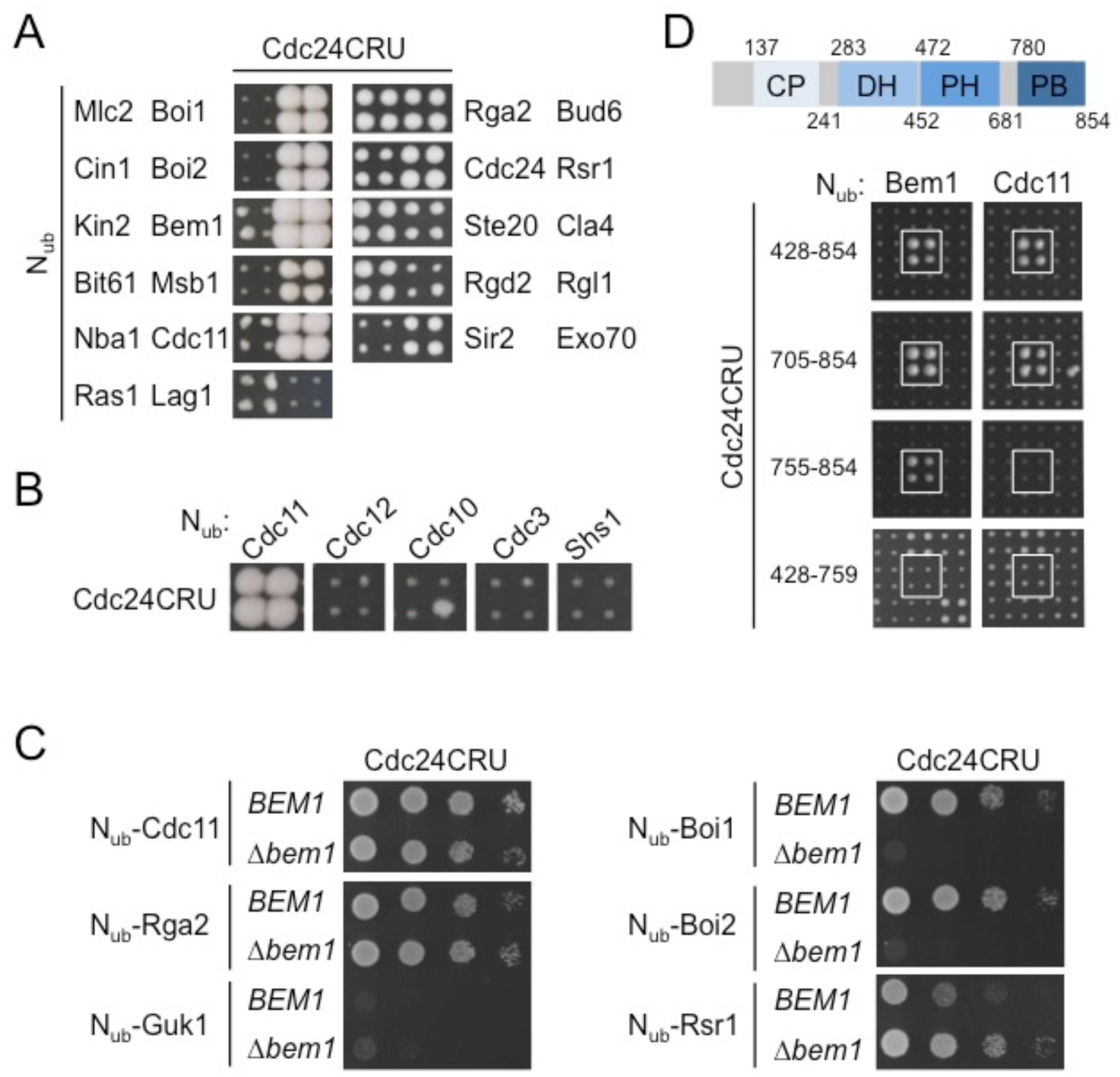
Cdc24 interacts with the septin subunit Cdc11. (A) Cut outs of a Split-Ubiquitin array, displaying diploid cells each co-expressing Cdc24CRU together with a certain N_ub_ fusion. Growth of a quadruplet of four independent matings on media containing 5-FOA indicates interaction between fusion proteins. N_ub_ fusions were expressed from the un-induced *P_CUP1_* promoter, except the N_ub_ fusions to Rga2, Bud6, Cdc24, Rsr1, Ste20 and Cla4 that were expressed from their native promoters. (B) Same as in (A) but showing cells expressing the N_ub_ fusions to the five mitotic septins under control of the *P_CUP1_* promoter. (C) Split-Ubiquitin assay of cells lacking either *BEM3,* or *BEM3* and *BEM1* and co-expressing Cdc24CRU together with the indicated N_ub_ fusions. Cultures were grown to OD_600_=1 and four µl of the undiluted or of 10-fold serial dilutions were spotted on media containing 5-FOA. (D) Upper panel: Domain structure of Cdc24: Numbers indicate the prospective first and last residue of the respective domains. Lower panel: as in (A) but with cells expressing CRU fusions to fragments of *CDC24* from centromeric plasmids. Boxed are the quadruplets of cells co-expressing N_ub_-Bem1 or N_ub_-Cdc11.

**Table 1.** List of Cdc24 interaction partners. List of N_ub_ labeled interaction partners of Cdc24CRU after subtraction of known false positives and chaperones as defined by Hruby et al. (Hruby et al., 2011), and updated in Figure S1. Complete list of arrayed N_ub_ fusions are shown in Figure S1.

|  | Cdc24-CRU |
| --- | --- |
| N<sub>ub</sub> fusion proteins | Bem1 |
|  | Boi1 |
|  | Boi2 |
|  | Bud6 |
|  | Cdc11 |
|  | Cdc24 |
|  | Cla4 |
|  | Exo70 |
|  | Msb1 |
|  | Ras1 |
|  | Rga2 |
|  | Rgd2 |
|  | Rsr1 |
|  | Ste20 |

To localize the binding site for Cdc11 on Cdc24 we screened fragments of Cdc24 as CRU fusions against N_ub_-Cdc11. A C-terminal fragment (Cdc24_428-854_) harboring the PH and PB domain of Cdc24 seem to contain the intact interface to Cdc11 and Bem1 (Fig. 1D). Deletion of the C-terminal PB domain in Cdc24_420-759_ abolished the interaction to Bem1 but also to Cdc11. Reversely, N_ub_-Bem1 but not N_ub_-Cdc11 generated an interaction signal with Cdc24_755-854_CRU (Fig. 1D). We conclude that PB_Cdc24_ provides the complete interface for Bem1 whereas the binding site for Cdc11 seems to further extend from the PB-towards the PH-domain of Cdc24.

A well-described mutation in the PB domain of Cdc24 that interferes with the formation of the Cdc24-Bem1 complex (Cdc24_428-854D820A_) does not affect the binding to Cdc11 (Fig. 2A) (Yoshinaga et al., 2003). This might indicate that Bem1 and Cdc11 contact different interfaces of PB_Cdc24_. To test whether Cdc24 can simultaneously bind Cdc11 and Bem1, we reconstituted the complex *in vitro*. *E. coli*-expressed GST-Cdc11 precipitated the PB domain of Bem1 specifically and only in the presence of Cdc24_420-854_ (Fig. 2B), thus proving the existence of a trimeric Cdc11-Cdc24-Bem1 complex. As cellular Cdc11 is predominately found in septin rods, we compared the interaction of 6His-Cdc24_420-854_ with Cdc11 in its free and its rod-incorporated state. Surface plasmon resonance spectrometry (SPR) showed that Cdc11 and the septin rod bind with comparable affinity to Cdc24 (Fig. 2C).

**Figure 2.**
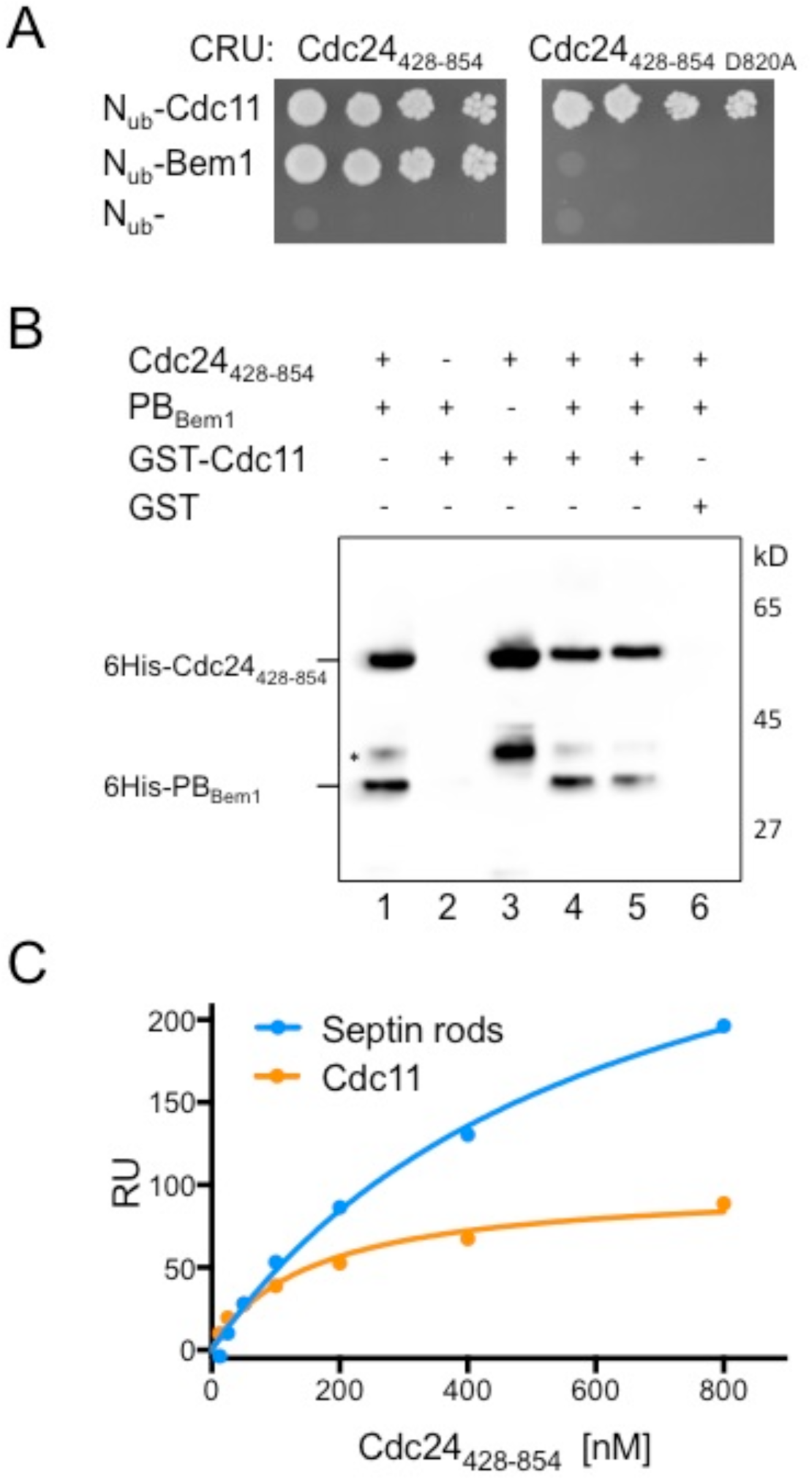
A trimeric complex of Bem1, Cdc24, and Cdc11. (A) Split-Ub assay as in Fig. 1C but of cells co-expressing the indicated N_ub_ fusion proteins and Cdc24_428-854_CRU (left panel), or Cdc24_428-854D820A_, containing a mutation in the binding interface for Bem1 (right panel). (B) GST-Cdc11-(lanes 2-5) or GST-coupled beads (lane 6) were incubated with 6His-PB_Bem1_-SNAP (lane 2), or 6His-Cdc24_428-854_ (lane 3), or 6His-Cdc24_428-854_ together with increasing concentrations of 6His-PB_Bem1_-SNAP (lanes 4, 5). Lane 1 shows the mixture of 6His-PB_Bem1_-SNAP and 6His-Cdc24_428-854_ before addition of beads. Glutathione eluates (lanes 2-6) were separated by SDS-PAGE and stained with anti-His antibody after transfer onto nitrocellulose. * indicates a proteolysis product of 6His-Cdc24_428-854_. Figure S2 shows the corresponding Coomassie staining of the gel. (C) Cdc11 or entire septin rods were immobilized on SPR-chips and incubated with increasing concentrations of purified 6His-Cdc24_428-854_. Binding of 6His-Cdc24_428-854_ to the chip-coupled septins is given in response units (RU). The K_D_s of the Cdc24_428-854_ - Cdc11 complex and the Cdc24_428-854_-septin rod complex were calculated to be 257 nM (± 47 nM; n= 4) and 351 nM (± 178nM; n=3) respectively.

### Cdc24 binds septins exclusively in G1 during bud site assembly

Live cell imaging of cells co-expressing Cdc11-mCherry and Cdc24-GFP restricted co-localization of both proteins to the prospective bud site shortly before bud emergence (PBS), and to the bud neck during cytokinesis (Fig. 3A). We monitored the interaction between Cdc11 and Cdc24 through the cell cycle by SPLIFF-analysis to resolve the interaction between both proteins in time and space. SPLIFF is a modification of the Split-Ub technique where the C_ub_ is sandwiched between the auto-fluorescent mCherry and GFP (CCG) (Moreno et al., 2013). Upon interaction-induced reassociation with a N_ub_ fusion, the GFP is cleaved off and rapidly degraded. The subsequent local increase in the ratio of red to green fluorescence indicates where and when the interaction between both proteins took place. Haploid yeast cells co-expressing Cdc11-mCherry-C_ub_-GFP (Cdc11CCG) together with N_ub_-Cdc24 were observed by fluorescence microscopy in intervals of 2 min. The N_ub_-induced conversion of Cdc11CCG to Cdc11CC was plotted as ratio of conversion against time (Fig. 3B). Interaction is indicated by a rise in conversion whereas a decreased or constant ratio indicates no interaction (Moreno et al., 2013). Accordingly, the analysis detects a clear interaction between Cdc11 and Cdc24 at the PBS that lasts for approximately four minutes during the assembly of the septins in the G1 phase of the cell cycle (Fig. 3B). No interactions between Cdc11 and Cdc24 were detected at the septin ring, the septin collar or the split-septin structure during later stages of the cell cycle (Fig 3B). A close inspection of the still images shortly before and during the assembly of the new septin patch at the PBS revealed that the labeled Cdc11 appears at the PBS predominantly in its converted Cdc11CC form (red fluorescence), whereas the fraction of Cdc11 at the old division site of mother and daughter cell consists predominantly of the unconverted Cdc11CCG (Fig. 3C, D). The assay does not resolve whether Cdc11CCG is converted by N_ub_-Cdc24 on flight to the PBS or immediately upon arrival. The latter being the more likely interpretation as the Cdc24-Bem1 complex arrives at the PBS shortly before Cdc11 (Okada et al., 2013; Lai et al., 2018) (see also Fig. 5)

**Figure 3.**
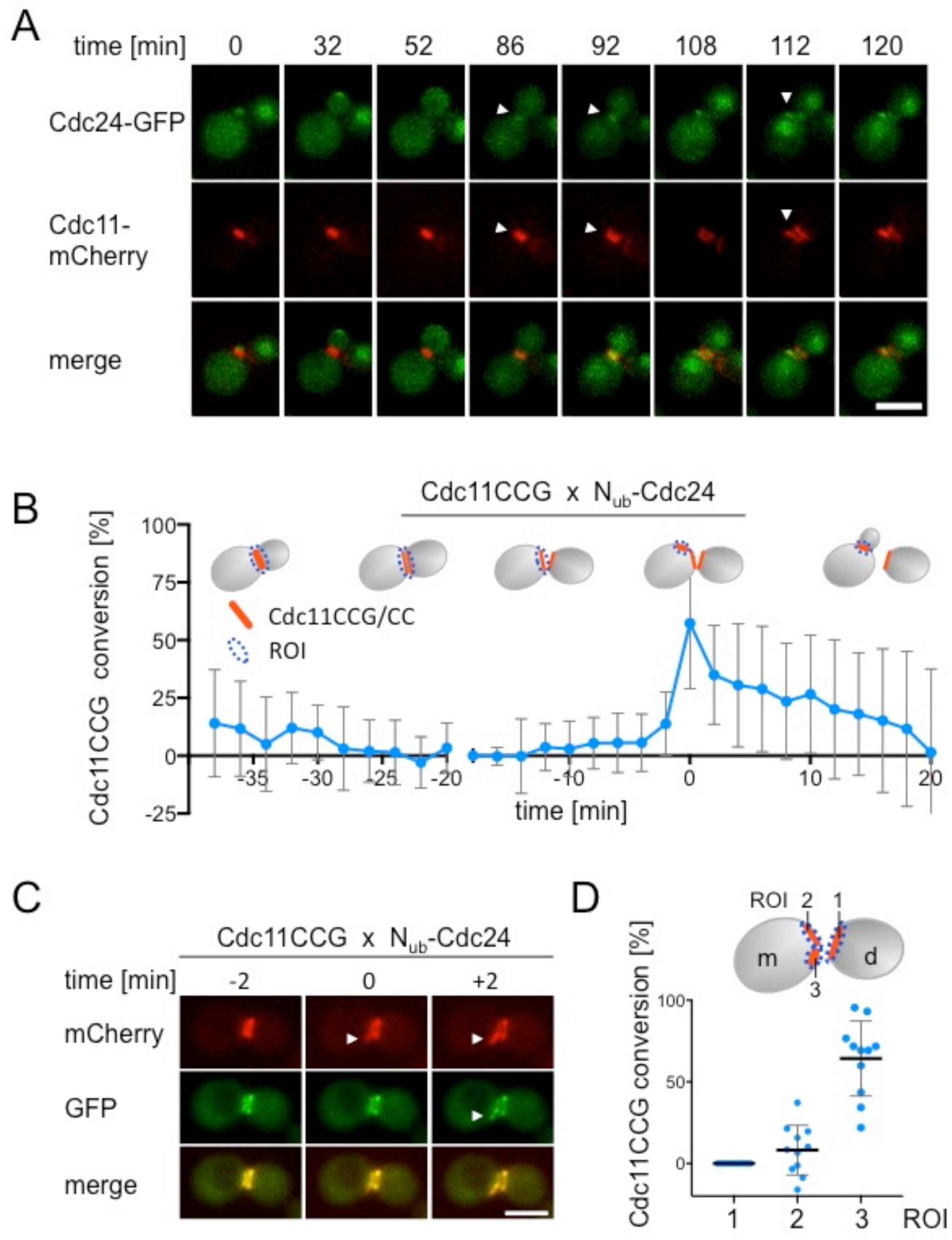
Cdc24 interacts with Cdc11 during G1 shortly before bud growth. (A) Time-lapse analysis of cells co-expressing Cdc24-GFP and Cdc11-mCherry. Arrowheads indicate co-localization occurring during cytokinesis at the bud neck (min 86, 92) and in G1 during PBS assembly (min 112). (B) SPLIFF-analysis of haploid cells co-expressing Cdc11CCG and N_ub_-Cdc24. The fraction of converted Cdc11CCG is plotted against time (n=11; SEM). The cartoon of the yeast cells indicates the cell cycle stage and the quantified region of the cell (ROI= region of interest). (C) SPLIFF analysis of N_ub_-Cdc24-induced conversion of Cdc11CCG during PBS assembly in the mother cell. Arrowhead indicates the site of bud assembly. Note the exclusive mCherry-staining at the PBS at min 0 (arrowhead in the middle panel). (D) Quantification of converted Cdc11CCG at min 0 (as defined in (C)) at the three indicated positions (ROI1-3) (n= 11; SEM). Significant conversion takes only place at the PBS of the mother cell (ROI3) but not at the sites of previous cell separation (ROI1, ROI2). Scale bar: 5µm.

### Binding to Cdc11 does not influence the GEF activity of Cdc24

Cdc11 and Cdc24 are essential proteins. To analyze the functional significance of the Cdc11-Cdc24-Bem1 complex we searched for mutations in Cdc24 that specifically impair the interaction with Cdc11. Cdc24 requires its PB domain for interacting with Cdc11 (Fig. 1D). The demonstration of a trimeric Cdc11-Cdc24-Bem1 complex implies that the contact areas of Cdc11 and Bem1 on Cdc24 do not overlap (Fig. 2). We thus exchanged a conserved Lysine against an Alanine at position 801 within the PB domain of 6His-Cdc24_428-854_, opposite the binding interface for Bem1. SPR analysis of the Cdc11-6His-Cdc24_428-854 K801A_ complex showed a modest reduction in binding affinity (Fig. 4A). Cdc11 also binds to regions of Cdc24_428-854_ that are N-terminal to its PB domain (Fig. 1D). We searched for conserved, exposed residues within Cdc24_428-854_ and exchanged the conserved Lysine in position 525 of the PH domain of Cdc24 against a glutamic acid (Cdc24_428-854 K525E_). SPR analysis of this mutant revealed a reduced affinity to Cdc11 (Fig. 4A) and a faster dissociation form the immobilized Cdc11 (Fig. 4A). Combining both mutations in one molecule (Cdc24_428-854 K525E K801A_, Cdc24_428-854 KK_) further decreased the affinity towards Cdc11 (Fig. 4A; equilibrium still not reached at 1.6 µM of analyte) and, compared to Cdc24_428-854_, increased the off-rate from the chip (Fig. 4B).

**Figure 4.**
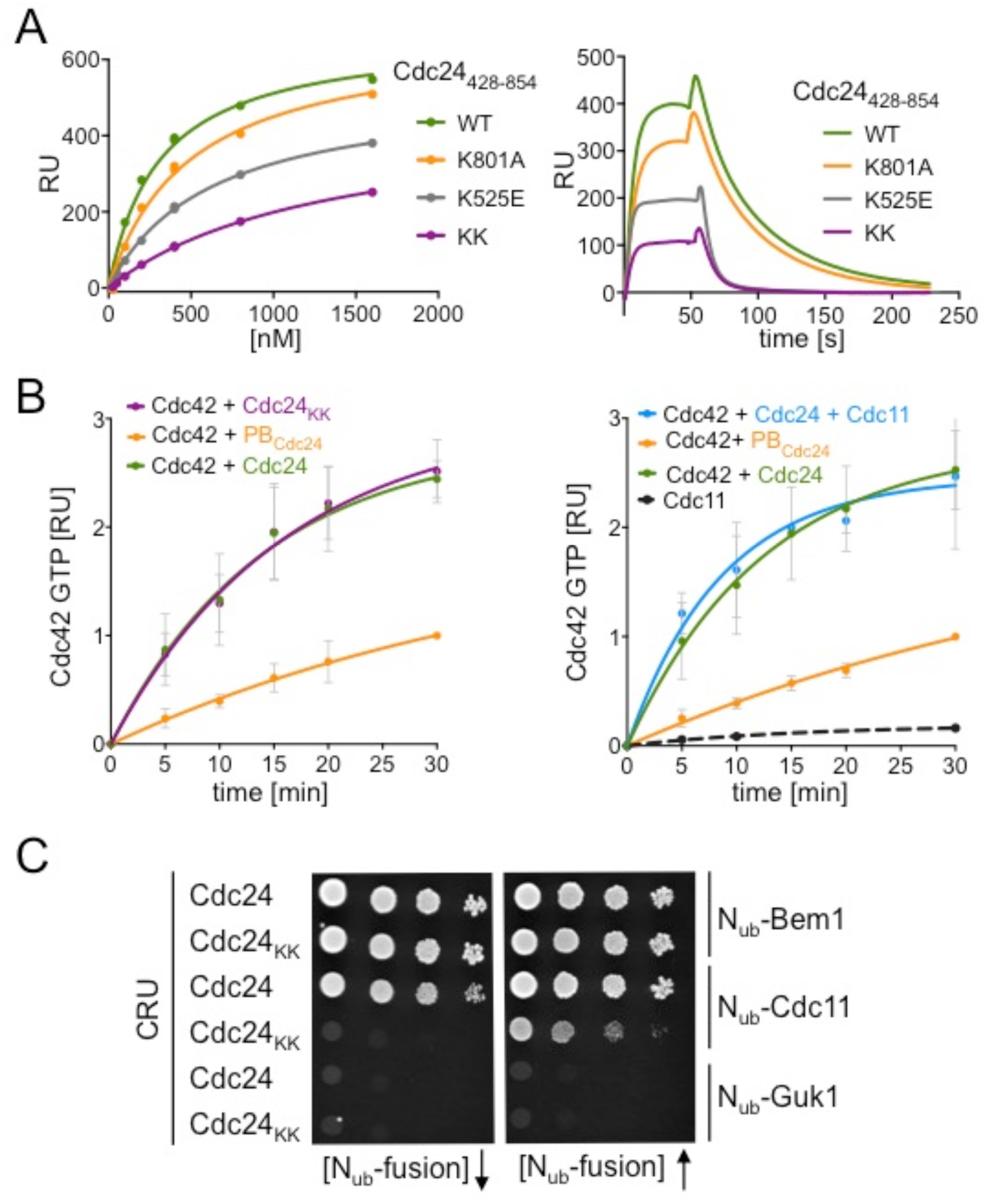
A mutant of Cdc24 with reduced affinity to Cdc11 but unaltered enzymatic activity. (A) Left panel: Cdc11 was coupled to SPR-chips and incubated with increasing concentrations of Cdc24_428-854_ carrying no, the single, or the double mutations. Shown are the increase in mass (RU) upon addition of the same amounts Cdc24_428-854_ and its mutants. Right panel: Sensograms of the 400 nM values of the right panel. The analyte solution was kept in the flow chamber for 60 s, before being washed for 180 s with analyte-free buffer. Note the nearly instantaneous loss of Cdc24_428-854KK_ from the Cdc11-coated chip after start of the washing step. (B) Enzymatic activities of Cdc24 and its mutant Cdc24_KK._ Left panel: Purified Cdc42 was incubated for 30 min with alpha-P^32^ GTP and MBP fusions to Cdc24, Cdc24_KK_, or the GST fusion to PB_Cdc24_, all enriched from bacterial lysates. Sample were taken at the indicated times and the amount of protein-bound alpha-P^32^ GTP was determined (n= 3; SD). Right panel: As in left panel but in addition showing the activity of Cdc24 in the presence of 1.6 µM Cdc11, and as control the binding activity of Cdc11 towards alpha-P^32^ GTP (n= 4; SD). (C) Cells expressing genomically integrated CRU fusions of *CDC24* or *cdc24_KK_* and the indicated N_ub_ fusions were spotted in tenfold serial dilutions onto media containing 5-FOA (left panel) or 5-FOA and 50µM of copper sulfate to induce the expression of the N_ub_ fusions (right panel). Note the light interaction signal between Cdc24_KK_CRU and N_ub_-Cdc11 upon N_ub_-Cdc11 overexpression.

Full length Cdc24 and Cdc24_KK_ were expressed and enriched as MBP fusions from *E. coli*-cells, or as TAP-tagged fusion from yeast cells (Fig. S3). Wild type and mutated forms of Cdc24 displayed in both instances roughly the same GDP/GTP exchange activity towards Cdc42 (Fig. 4B, Fig. S4). The addition of purified 6His-Cdc11 in excess over MBP-Cdc24 did not significantly alter the activity of Cdc24 either (Fig. 4B). We conclude that neither the interaction-impairing mutations nor the binding to Cdc11 seem to affect the catalytic activity of Cdc24.

We introduced the two interaction-impairing mutations into the genomic sequence of *CDC24* to create the allele *cdc24_KK_*(*cdc24_K525E K801A_*) in yeast. In accordance with our *in vitro* analysis, a CRU fusion to *cdc24_KK_* monitored a strongly impaired but not completely abolished interaction with N_ub_-Cdc11 (Fig. 4C). Cdc24_KK_CRU retained its binding to N_ub_-Bem1 (Fig. 4C). Cells containing the *cdc24_KK_* allele did not display any obvious growth phenotype (see also Fig. 6C).

### The Cdc11-Cdc24-Bem1 complex promotes the formation of the PBS

Cdc24_KK_-GFP, or Cdc11-mCherry and Bem1-GFP in *cdc24_KK_*-cells displayed a wild type-like distribution (Fig. 5A, B). However, a closer inspection of the cell cycle phase in which both proteins interact revealed that Cdc24_KK_-GFP is not always permanently fixed to a once chosen PBS (Fig. 5A, Movies S1, S2). During G1 the Cdc24_KK_-GFP patch, indicative of the PBS, occasionally dissolved and reappeared at the site of previous cell separation or less often at an entirely new site of the plasma membrane (Fig. 5A, B, Movie S2). Bem1-GFP, or Gic2_PBD_-RFP, a visual reporter of Cdc42_GTP_, showed a similar retraction from the initially chosen site of bud formation in *cdc24_KK_*-cells (Fig. 5B) (Brown et al., 1997; Atkins et al., 2013; Orlando et al., 2008).

**Figure 5.**
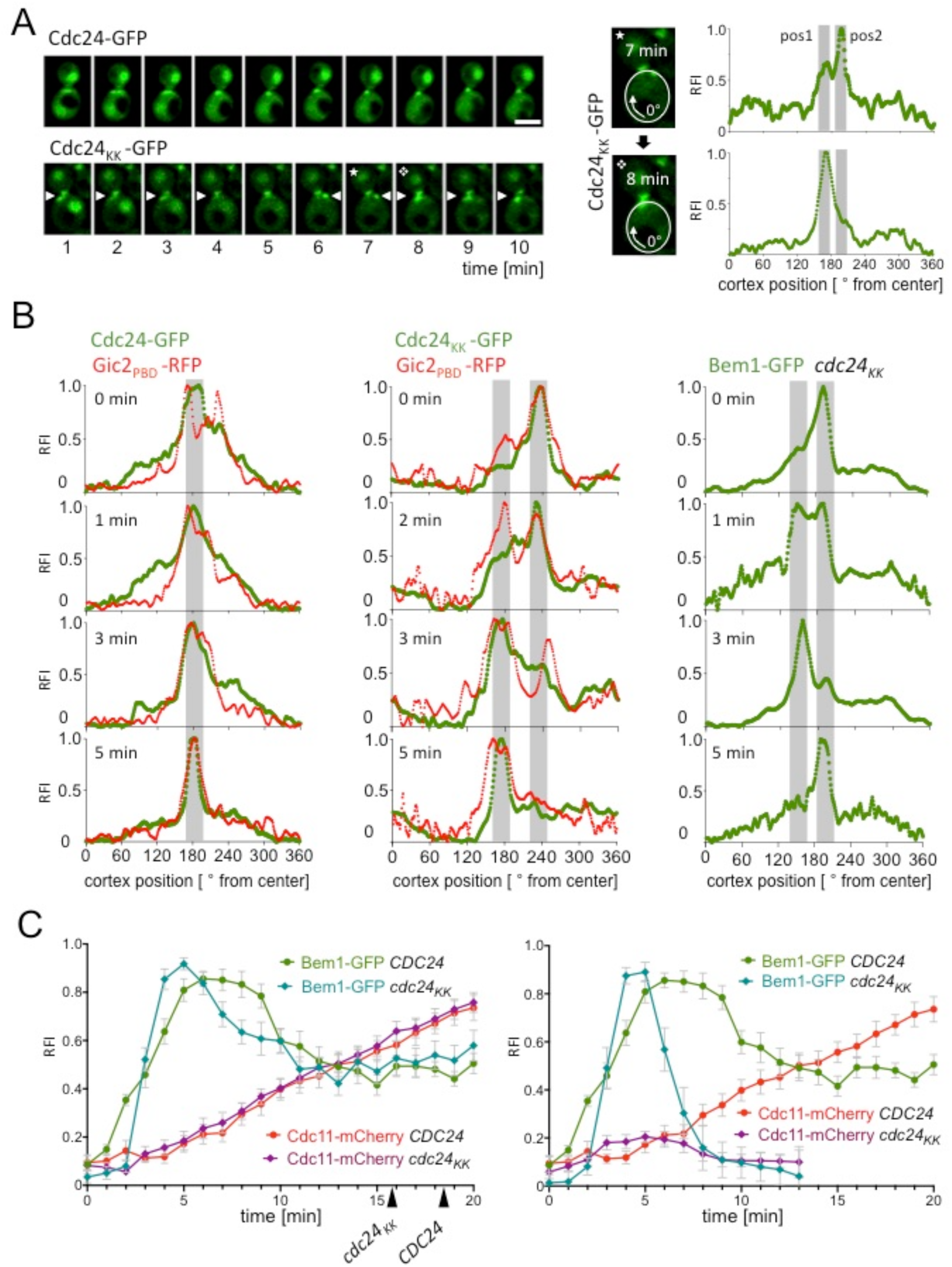
The interaction between Cdc24 and the septins enforce PBS stability and septin recruitment. (A) Left panel: GFP fusions to Cdc24 or Cdc24_KK_ were recorded by time-lapse microscopy over time. Note the changing position of Cdc24_KK_-GFP in the mother cell between min 7 and 8. Right panel: Blow up of min 7 and 8 of left panel and the quantifications of the GFP intensities at the given positions of the cortex. Scale bar: 5 µm. (B) Fluorescence intensity profiles of the cortex of cells expressing Cdc24-GFP and Gic2_PBD_-RFP (left panel), Cdc24_KK_-GFP and Gic2_PBD_-RFP (middle panel), or Bem1-GFP in *cdc24_KK_*-cells (right panel). Profiles were measured in short time intervals during PBS formation of mother cells as in (A). (C) Time-lapse analysis of wild type- or *cdc24_KK_*-cells co-expressing Bem1-GFP and Cdc11-mCherry during PBS formation in mother cells. Left panel: Intensity profiles form wild type cells (n=13) and the fraction of *cdc24_KK_*-cells (n=13) that keep the original position of the PBS. Right panel: Intensity profiles from the fraction of *cdc24_KK_*-cells (n=13) that retract from the position of the original PBS. The profile of the wild type cells from the left panel is given as reference. All error bars indicate SEM. The arrows indicate the first visible separation between GFP- and mCherry fluorescence and thus the start of bud growth.

We next visualized the dynamics of the Cdc24-Bem1 complex together with the septins during PBS formation in *cdc24_KK_*- and wild type cells (Fig. 5C). The observed wild type cells co-expressing Cdc11-mCherry and Bem1-GFP behaved uniformly. Bem1-GFP rapidly accumulated, kept its high intensity at the PBS for five minutes before trailing to approximately 50% of its peak intensity in the next 4 min (Fig. 5C). A decrease in intensity of Bem1 during PBS formation was already reported and was also observed for active Cdc42 (Woods et al., 2016; Okada et al., 2013). Starting shortly after the initial rise in Bem1-GFP fluorescence, Cdc11-mCherry accumulated steadily, and reached approximately 50% of its final intensity at the PBS when the signal of Bem1-GFP had already leveled off to 50% of its maximal intensity (Fig. 5C).

Bud site assembly occurred by two alternative ways in *cdc24_KK_*-cells (Fig. 5C). In 70% of the measured mutants the initially chosen PBS was kept, and a new bud was formed next to the previous division site (Fig. 5C). Here, Bem1-GFP intensity rapidly rose to similar levels as in wild type cells but stayed at this intensity for only two minutes. An initial rapid decrease was followed by a much slower decline that finally trailed off to 50% of the intensity of its peak value (Fig. 5C). The accumulation of Cdc11-mCherry occurred indistinguishable from wild type cells.

The remaining 30% of mutant cells abort bud site assembly (Fig. 5C). In these cells Bem1-GFP initially appeared with similar kinetics at the PBS, but stayed at its peak value for only approximately 2 min before rapidly dissolving (Fig. 5C). Cdc11-mCherry accumulation stopped early in these cells, and the small fraction of attached Cdc11 disappeared together with Bem1 from the PBS (Fig. 5C). Both proteins start a new round of PBS assembly, preferentially at the site of former cell division.

### Genetic analysis positions the Cdc11-Cdc24 interaction in the Cla4 assembly pathway

Bem1 is a major determinant for Cdc24 localization and a stimulator of its catalytic activity (Smith et al., 2013; Rapali et al., 2017; Woods et al., 2015). Cells carrying the *cdc24_KK_* allele do not tolerate the deletion of *BEM1* (Fig. 6A). This outcome disfavors the possibility that the two mutations in Cdc24_KK_ interfere with Bem1’s positive influence on Cdc24 but suggests instead that Cdc11 and Bem1 might cooperate to keep Cdc24 at the PBS, or that Cdc24 and Bem1 contribute independently to septin recruitment. To ask more directly whether the impaired interaction to Cdc11 causes the synthetic lethality between *cdc24_KK_* and *BEM1*, we fused the PB-domain of Bem1 to the C-terminus of Cdc11 (Cdc11-PB_Bem1_, Fig. 6B). Cdc11-PB_Bem1_ will connect Cdc11 to Cdc24 irrespective of the presence of the two Cdc11 interaction-impairing mutations in the *cdc24_KK_* allele. Fig. 6B shows that the ectopic expression of Cdc11-PB_Bem1_ rescues the growth of *cdc24_KK_* Δ*bem1-*cells (always in a *Δbem3* background), thus confirming the critical contribution of the Cdc24-Cdc11 interaction to bud site assembly.

To map the interaction between Cdc24 and Cdc11 within the genetic network of bud site assembly we compared the growth of strains carrying *cdc24_kk_* and the deletion of a certain member of the polarity genes. Cells lacking any of the tested polarisome subunits, or Rsr1 are strongly affected by *cdc24_kk_,* whereas cells carrying deletions of *CLA4,* or *STE20* are not (Fig. 6C). When *Δspa2 cdc24_kk_*-cells were shifted from 30°C to 37°C for four hours, the majority of cells arrested at large unbudded cells (Fig. 6D). The analysis positions the Cdc11-Cdc24 interaction parallel to the polarisome/actin pathway and within the Cla4 pathway of bud site- and septin ring assembly.

**Figure 6.**
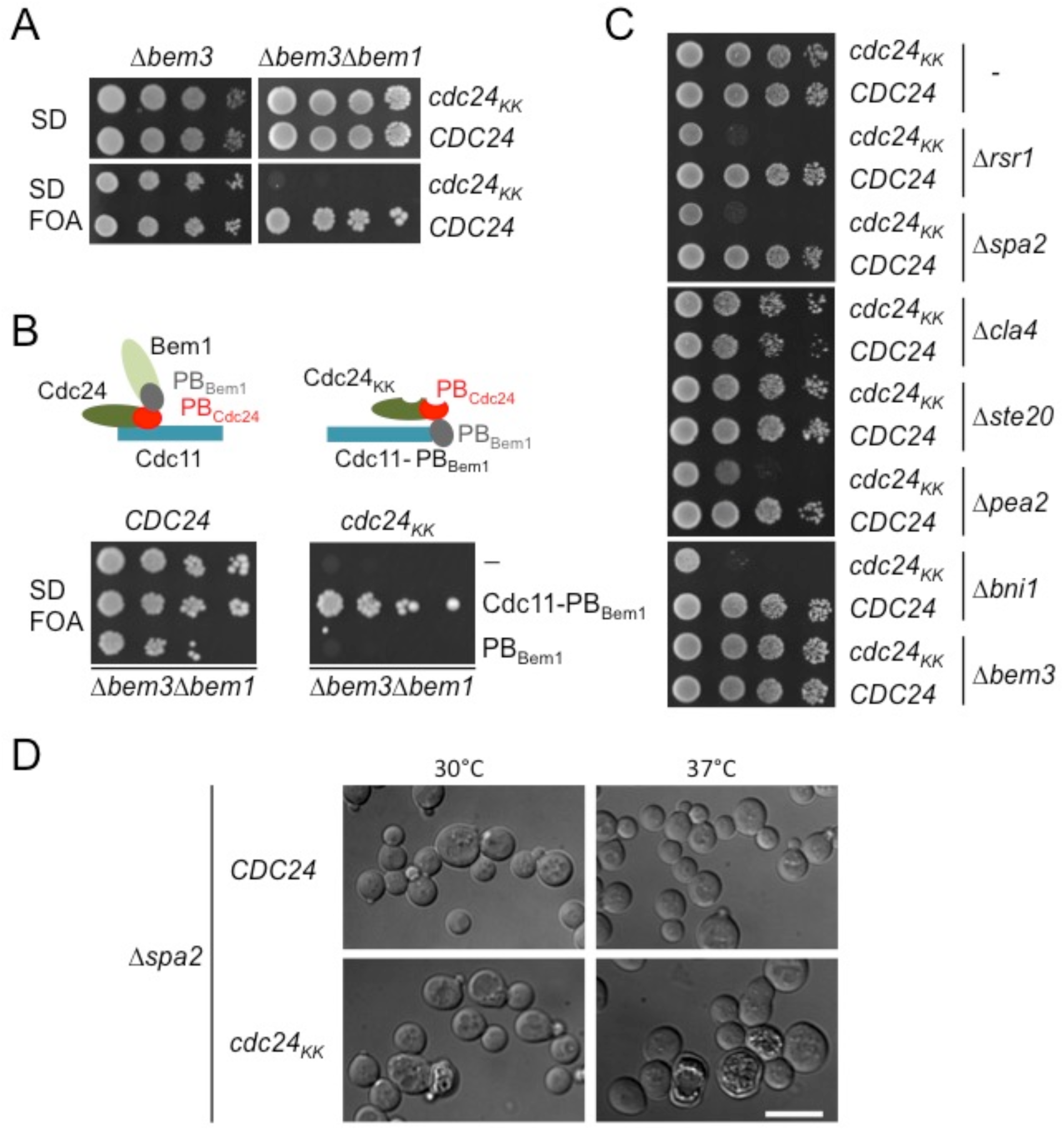
Genetic interactions of the *cdc24_KK_* allele. (A) Wild type cells or cells expressing *cdc24_kk_* and lacking *BEM3* (left panel), or lacking *BEM3* and *BEM1* (right panel), and containing an additional plasmid-based copy of *CDC24* were grown to an OD_600_ of 3 and spotted in 10-fold serial dilutions on SD, or SD plus 5-FOA to counter-select against the plasmid-encoded *CDC24.* (B) Upper panel: Cartoon of the Cdc24-Cdc11 complexes as possibly realized in the cells below. Left: Bem1-Cdc24-Cdc11 complex in wild type cells. Right: Cdc24_KK_ bound to an artificial chimera between Cdc11 and the PB domain of Bem1 in *Δbem1*-cells. Highlighted are the PB domains of Bem1 in grey and of Cdc24 in red. Lower panel: As in (A) but with *Δbem3Δbem1*-cells carrying in addition an empty plasmid, or a plasmid expressing Cdc11-PB_Bem1_, or PB_Bem1_ from a P*_MET17_* promoter under slightly repressing conditions (70µM methionine). (C) Cells of the indicated genotypes were grown to an OD_600_ of 1 and spotted in 10-fold serial dilutions on SD. Cells were incubated at 37 °C for three days. (D) *Δspa2*-cells either carrying *CDC24* or *cdc24_KK_* were shifted to 37 °C for 2,5 hours and observed by DIC microscopy. Scale bar: 10 µm.

### Impaired Cdc24-septin interaction increases budding through the former cell division site

Fluorescence microscopic analysis of *cdc24_KK_*-cells co-expressing Cdc11-mCherry and Bem1-GFP predict that approximately 18% of *cdc24_KK_*-cells should bud within the previous cell division site (Fig. 7A). The expected morphological consequences were visualized by comparing scanning electron microscopic pictures obtained from wild type and *cdc24_KK_* cells (Fig. 7A). For the quantitative comparison we included buds forming on the rim of an old bud- or birth scar as bud in scar phenotype. Whereas each single mutation had only a small effect on cellular morphology, *cdc24_KK_* cells showed a twofold higher incidence of bud in scar growth (35%) than wild type cells (18%) (Fig. 7A). The increase of 17% match the fraction of *cdc24_kk_*-cells cells that, according to fluorescence microscopy, relocate their PBS from its initial axial site back to the old division site (Fig. 7A). Consequently, buds that emerged centrally and repeatedly through a former bud- or birth scar were frequently observed in EM images of *cdc24_KK_* cells but never in wild type cells (Fig. 7A).

**Figure 7.**
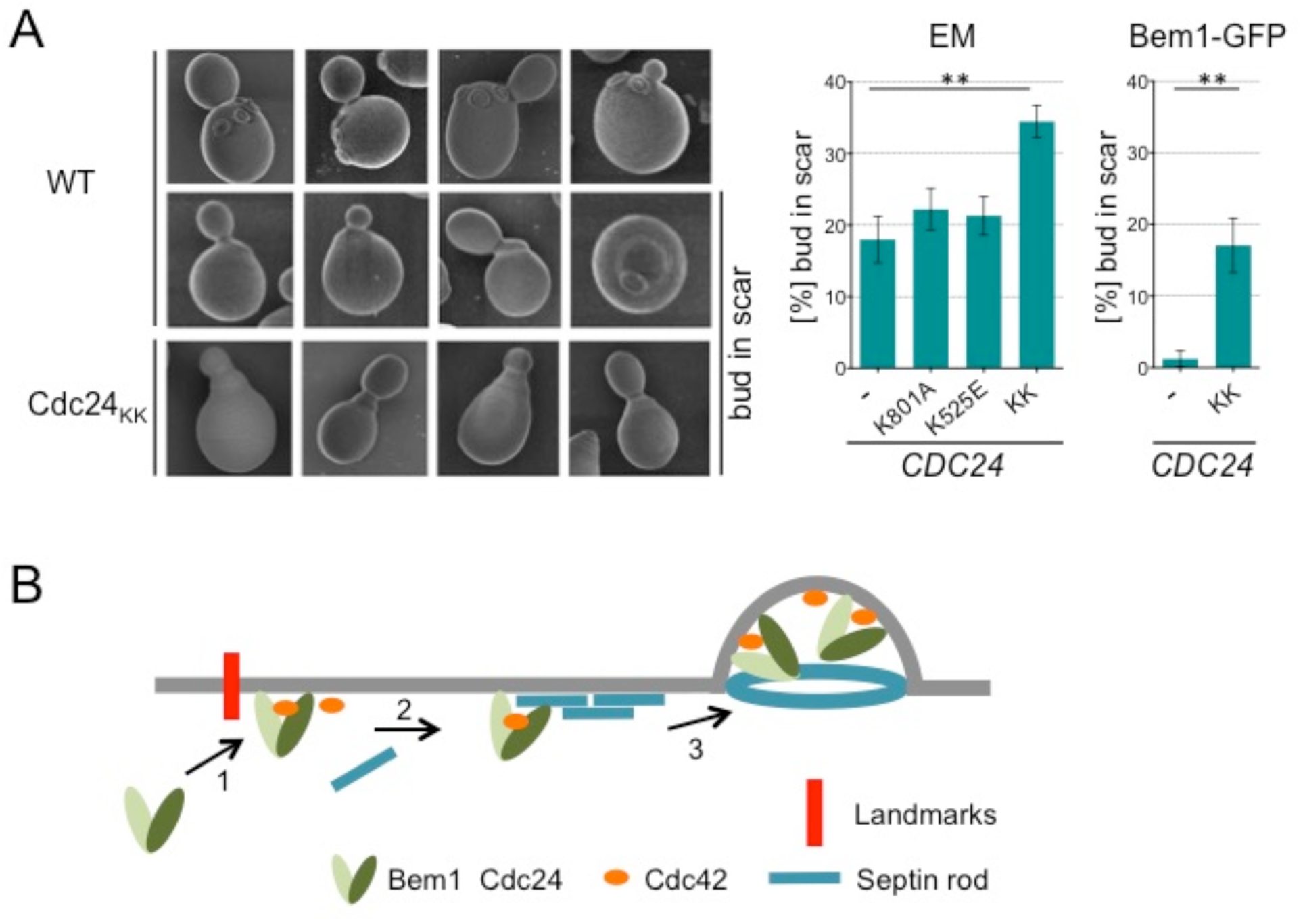
The impaired interaction between Cdc11 and Cdc24 increase budding in the old division site. (A) Left panel: EM pictures of wild type cells showing axial budding (upper row) and budding at the rim of a birth/bud scar (middle row), and of *cdc24_KK_-*cells showing repeated budding through a previous bud site (lower row). Middle panel: Quantification of the budding phenotypes of cells carrying the indicated alleles of *CDC24*. Fraction of cells with “bud in scar” phenotype: *CDC24*: 18,0% (SD ± 3.3; n=569). *cdc24_K801A_*: 22,2% (SD ± 2.9; n=552). *cdc24_K525E_*: 21,3% (SD ± 2.7; n=489). *cdc24_KK_*: 34.5% (SD ± 2.3; n= 450). ** P value= 0.002. Right panel: The extent of budding through the previous bud site as quantified by fluorescence microscopy of *CDC24-* (n= 158) or *cdc24_KK_*-cells (n= 151) expressing Bem1-GFP (Error bar SEM; ** P= 0.0024). (B) Model of the function of the Cdc24-Cdc11 complex. (1) Landmark proteins including Rsr1 attract and activate the Bem1-Cdc24 complex. Ongoing production of Cdc42_GTP_ recruits polarity proteins. (2) Cdc42_GTP_ and the Cdc24-Bem1 complex attract the septins to the PBS. The septins tighten the association of Cdc24-Bem1 complex at the PBS. If septin recruitment fails Bem1-Cdc24 leaves the PBS to start the assembly process again. (3) Ring formation of the septins displaces the Bem1-Cdc24 complex to center of the ring to initiate the outgrowth of the bud.

## Discussion

The Cdc24-Bem1 complex generates active Cdc42 that is thought to recruit the septins and other effectors to the PBS. The discovery of a direct interaction between Cdc24 and Cdc11, the capping subunit of the septin rod, points to a more direct role of the Cdc24-Bem1 complex in septin recruitment and PBS assembly. The Cdc24-Cdc11 interaction is under strict cell cycle control and occurs within 4 min during the formation of the PBS. Timing and location of the interaction suggest that the Cdc24-Bem1 complex might help to directly recruit the septins. However a mutant of Cdc24 with significantly reduced affinity to Cdc11 reveals a more complex role of the Cdc11-Cdc24 interaction during this process. Cells carrying the *cdc24_KK_* allele will either keep the once chosen site of bud formation or abort it and search for a new PBS. In the former cells, septin recruitment occurs without delay and wild-type speed. However, the Cdc24-Bem1 complex is less stably fixed to the PBS and a fraction of it leaves the PBS earlier than in wild type cells. This outcome argues for a role of the Cdc11-Cdc24 interaction in keeping the Cdc24-Bem1 complex at the PBS. The molecular choreography is different in *cdc24_KK_*-cells that abort bud site assembly altogether. Although Cdc24-Bem1 appears at the PBS with similar kinetics, only a small amount of septins gets recruited before the Cdc24-Bem1 patch rapidly dissolves. We consider the two alternative phenotypic manifestation of the *cdc24_KK_* allele as evidence for two successive and reinforcing roles of the Cdc11-Cdc24 interaction. In a first step it helps to attract the septins to the PBS, whereas in the second step it retains the Cdc24-Bem1 complex at the PBS (Fig. 7B).

Why is the additional retention of Cdc24-Bem1 complex needed? Wild type cells maintain the high concentrations of Cdc24-Bem1 complexes at the PBS for only four minutes before half of the complexes leave this site. Bem1 is mainly responsible for anchoring the complex through a mix of ligands including the Boi1/2 proteins, acid phospholipids and Cdc42_GTP_ (Bose et al., 2001; Kozubowski et al., 2008; Meca et al., 2019; Fairn et al., 2011). These initial core components then attract a large number of other proteins that might start to compete with Cdc24-Bem1 for limiting binding sites at the cortex. To avoid a complete displacement of Cdc24-Bem1, an additional stabilization of Cdc24-Bem1 might be provided by the attracted septins through a novel interface not recognized by other polarity proteins. As the enzymatic activity and the direct interaction to Cdc24 originally guides the septins to the PBS, the mechanism is best described as a positive feedback between Cdc24 activity and septin recruitment. This positive feedback rewards a polarity patch that has successfully attracted the septins, and enforces a linear order of its assembly (Fig. 7B).

The recruitment of the septins to the PBS occurs slightly later than the majority of many other polarity proteins. This delay can be explained by the necessity to prime the septins or their receptor by either CDK or a CDK-activated mediator (Lai et al., 2018). Our results suggest the Cdc11-Cdc24 interaction as a candidate for this priming event. Phosphorylation of either of the two components might increase their affinity to each other and induce the assembly of the septins at the PBS. Cla4 is known to bind Bem1 and to phosphorylate Cdc24 and the septins (Gulli et al., 2000; Versele and Thorner, 2004). In contrast to other polarity genes, a deletion of *CLA4* shows no negative genetic interaction with the *cdc24_KK_* allele. Cdc11-Cdc24 complex formation and Cla4-induced phosphorylations might thus be successive steps in the same septin assembly- and transformation pathway. Whether Cla4 serves as mediator of CDK and directly primes the interaction between Cdc24 and the septins is an open but experimentally testable question.

The Cdc24-Bem1 complex does not bind mature septin rings during later cell cycle stages (Fig. 7B). The dislocation of the Cdc24-Bem1 complex from the septins migh thus correlate with septin ring formation and might mark the transition between PBS assembly and incorporation of new plasma membrane and cell wall material. We favor the hypothesis that the release of Cdc24-Bem1 from the septins serves as a checkpoint that ensures that tip growth only begins once the septin patch has successfully transformed into a ring (Fig. 7B).

Previous experiments have shown that septins also exert a negative influence on polarity patch formation (Okada et al., 2013; Schneider et al., 2013). To reconcile both observations we postulate that a switch in its regulatory properties accompanies the transformation from septin rods to filaments and rings. Whereas rods bind the Cdc24-Bem1 complex and promote PBS formation, rings or filaments might exclude the Cdc24-Bem1 complex and recruit Cdc42 GAPs instead to inhibit or terminate PBS formation (Okada et al., 2013). The switch thus closely couples the maturation of the PBS with the structural transformations of septins. Both feedbacks are compatible with the growth of the septin structures before and after the switch. Whereas the initial septin recruitment depends on Cdc42_GTP_, the further assembly of the septins during and after the transformation to ring and collar is template-assisted and independent of Cdc42. Similar feedback mechanisms might operate during the assembly of higher-order septin structures in other organisms as well (Nagata and Inagaki, 2005).

## Material and Methods

### Growth conditions and cultivation of yeast strains

All yeast strains were derivatives of JD47, a descendant from a cross of the strains YPH500 and BBY45 (Dohmen et al., 1995). Cultivation of yeast was performed in standard SD or YPD media at 30°C or the indicated temperatures as described (Kustermann et al., 2017). Media for Split-Ubiquitin interaction assay and selection for the loss of centromeric *URA3*-containing plasmids comprised 1 mg/ml 5-fluoro-orotic acid (5-FOA, Formedium, Hunstanton, UK).

### Construction of plasmids, gene fusions and manipulations

Construction of N_ub_ and C_ub_ gene fusions as well as GFP-, mCherry- or mCherry-C_ub_-RGFP (CCG)-fusions was as described (Wittke et al., 1999; Dünkler et al., 2012; Neller et al., 2015; Moreno et al., 2013). Cdc24CRU/ -GFP, Bem1-GFP, Shs1-mCherry and Cdc11-mCherry/-CCG were constructed by genomic in-frame insertions of the *GFP-*, *CRU-*, or *CCG*-modules behind the coding sequences of the relevant genes. In brief, a PCR-fragment of the C-terminal region of the respective target gene lacking the stop codon was cloned via *Eag*I and *Sal*I restriction sites in front of the CRU-, GFP-, mCherry-, or CCG-module on a *pRS303*, *pRS304* or *pRS306* vector (Sikorski and Hieter, 1989). Plasmids were linearized using unique restriction sites within this sequence and transformed into yeast cells for integration into the genomic target ORF. Colony PCR with diagnostic primer combinations was used to verify the successful integration. *CDC24_428-854_*-*, CDC24_705-854_-, CDC24_755-854_*-, and *CDC24_428-759_CRU* were obtained by ligation of *Eag*I/*Sal*I-cut PCR fragments spanning the respective ORF between the sequences of the *P_MET17_* promoter and the CRU module on the *pRS313* vector (Sikorski and Hieter, 1989). Mutations in the coding region of *CDC24* were obtained by overlap-extension PCR using plasmids containing CDC24_428-854_ as templates.

Construction of genomic *cdc24_K525E_*, *cdc24_K801A_*, *cdc24_K525E K801A_* (*cdc24_KK_*), and *CDC24_WT_* alleles was achieved by recombination of PCR fragments generated from plasmids pFA6a*-Cdc24_428-854 KK_+term,* and pFA6a*-Cdc24_428-854_+term* using a forward primer starting at 428 aa and a standard reverse S2 primer annealing to a stretch behind base pair 186 of the *CDC24* terminator. C-terminal GFP fusions to *cdc24_KK_* and *CDC24_WT_* were created by PCR-based in-frame fusion using pYM26 and pYM28 as templates (Janke et al., 2004). P*_MET17_*-CDC11-PB_Bem1_ pRS313 was obtained by fusing a PCR product of the PB domain starting from residue 470 in frame of *CDC11* in a P*_MET17_* pRS313 vector. Cdc11 and PB_Bem1_ were separated by a linker sequence encoding a single HA epitope tag.

In certain strains the native promoter sequence was replaced by *P_MET17_* through recombination with a PCR fragment generated from pYM-N35 and primers containing sequences identical to the respective genomic locations at their 5’ ends (Janke et al., 2004). *GST* fusions were obtained by placing the ORF of the respective gene or gene fragment in frame behind the *E. coli GST* sequence on the pGEX-2T plasmid (GE Healthcare, Freiburg, Germany) using *Bam*HI and *Eco*RI restriction sites. Fusions to the SNAP-tag (New England Biolabs, Beverly, MA) were expressed from plasmid pAGT-Xpress, a pET-15b derivative (Schneider et al., 2013). Gene fragments were inserted in frame into a multi-cloning site located between the upstream *6HIS*-tag-coding sequence and the downstream *SNAP*-tag-coding sequence. The *6HIS*-tag fusions were obtained by placing the ORF of the respective gene or gene fragment behind the *E. coli 6HIS*-tag sequence on the previously constructed pAC plasmid (Schneider et al., 2013). Fusions to MBP were obtained by inserting the ORF of *CDC24* or its mutants in the multi-cloning site of vector pMAL-c5X (New England Biolabs, Beverly, MA).

Gene deletions were performed by one step PCR-based homologous recombination using pFA6a natNT2, pFA6a hphNT1, pFA6a kanMX6, pFA6a CmLEU2 and pFA6a HISMX6 as templates (Bähler et al., 1998; Longtine et al., 1998; Janke et al., 2004; Schaub et al., 2006). Lists of used and constructed plasmids and yeast strains can be found in Table S1 and S2.

### Split-Ubiquitin library screen

JD47 yeast cells expressing a genomic CRU fusion to *CDC24* were transformed with a multi-copy library expressing random fragments of the yeast genome inserted behind the N_ub_-module and driven by the *P_ADH1_* promoter (Laser et al., 2000). Library transformation, screening and the analysis of the FOA-resistant clones were as described (Reichel and Johnsson, 2005). A plasmid carrying a N_ub_ fusion to Cdc11 lacking the N-terminal five residues was independently isolated twice.

### Split-Ub interaction analysis

Split-Ubiquitin array analysis: A library of 548 different α-strains each expressing a different N_ub_ fusion were mated with a *CDC24*-*C_ub_-R-URA3* (CRU) expressing a-strain. Diploid were transferred as independent quadruplets on SD media containing 1 mg/ml 5-FOA and different concentrations of copper to adjust the expression of the N_ub_ fusions (Fig. S1) (Dünkler et al., 2012). Individual Split-Ub interaction analysis: CRU and N_ub_ expressing strains were mated or co-expressed in haploid cells and spotted onto medium containing 1 mg/ml FOA and different concentrations of copper in four 10-fold serial dilutions starting from OD_600_=1. Growth at 30°C was recorded every day for 3 to 5 days.

### Preparation of PB_Bem1_ affinity matrix

200 µl of ‘Glutathion Sepharose^TM^ 4 Fast Flow’ suspension (GE-Healthcare) was diluted in 500 µl of PBS and the beads were pelleted via centrifugation. After two additional washing steps with PBS, 2 ml of BL21-cell-lysate containing the fusion protein GST-PB_Bem1_ were added and rotated for 15 minutes at 4 °C. The beads were finally washed three times in PBS.

### Enrichment of Cdc24-TAP fusion

Yeast cells expressing Cdc24-TAP or Cdc24_KK_-TAP were grown in 500 ml of SD Trp-media at 30°C to an OD_600_ of 1.5, harvested by centrifugation, washed with H_2_O and shock frozen in liquid nitrogen. The frozen pellets were transferred to a pre-cooled mortar and grinded for at least 10 min to a fine powder under constant presence of liquid nitrogen. The powder was transferred to a new vessel, dissolved in 1 ml of ice cold lysis buffer (20 mM Tris base pH 7.5, 80 mM NaCl,1 mM DTT) containing a protease inhibitor cocktail (Roche Diagnostics, Penzberg, Germany). The solution was centrifuged for 10 min at 4 °C and 1600 g. 40 µl of magnetic hsIgG beads (Thermo Fisher Scientific, Waltham, MA, USA) were washed three times in lysis buffer and resuspended in the cleared yeast extract. After 4 h of overhead-rotation at 4 °C, the beads were thoroughly washed six times in 500 µl of 20 mM Tris base pH 8.0, 100 mM NaCl, 5 mM MgCl_2_, and used immediately for the GEF assay (without additional buffer).

### Enrichment of MBP-Cdc24 fusion

*E. coli* cells expressing either MBP-Cdc24 or MBP-Cdc24_KK_ were grown in 1L SB-Amp to an OD_600_ of 0.8, adjusted to 1 mM IPTG and cultivated at 18 °C over-night. The cells were harvested by centrifugation, washed with H_2_O and partitioned in 4 aliquots. Cell-pellets were resuspended in 25 ml PBS and incubated for 30 min at 4 °C in the presence of 1 mg/ml Lysozym and protease inhibitors (Roche Diagnostics, Penzberg, Germany) The cell lysate was cleared by centrifugation at 40000 g for 10 minutes after sonification with a Bandelin Sonapuls HD 2070 (Reichmann Industrieservice, Hagen, Germany). 25 ml of the MBP-Cdc24 extract was added to 200 µl PB_Bem1_ affinity matrix and the suspension was rotated for 2 h at 8 °C. The supernatant was removed, the beads were resuspended in PBS, transferred to a Mobicol “F” column (Mobitec GmbH Göttingen, Germany) and washed 3 times with PBS. To elute the bound protein, 250 µl of 10 mM reduced glutathione in 50 mM Tris pH 8.0 was added and the suspension was incubated for 10 minutes at 10°C. After centrifugation, the flow trough was transferred in 25 mM Tris base pH=8.0,100 mM NaCl, 5 mM MgCl_2_, 20% w/v Glycerol using a NAP5-column (GE Healthcare, Freiburg, Germany), partitioned in 10 µl aliquots and stored at -20°C for a maximum of 2 weeks.

### Cdc24 GEF assays

GEF activity of was quantified by recording the amount of [αP^32^]GTP (Hartmann Analytic, Braunschweig, Germany) taken up by 6His-Cdc42 in the presence of MBP-Cdc24, TAP-Cdc24 or 6His-Cdc24_760-854_ as control (Mionnet et al., 2008). 6His-Cdc42 was purified from crude *E. coli* extracts by IMAC and size exclusion chromatography. For each time-point, 40 pMol purified Cdc42 and 3 µCi [αP32]GTP were added to the assay-buffer (25 mM Tris base pH 8.0,100 mM NaCl, 5 mM MgCl2, 1 mM DTT, 1 mM EDTA, 5 µM GTP). After the addition of MBP-Cdc24 or MBP-Cdc24_KK_ the mixture was incubated under constant shaking at 22°C. 25 µl of the assay volume was removed at each time-point and quenched with 500 µl of ice-cold assay buffer. The samples were immediately filtered on pre-wetted 0.45 µm nitrocellulose filters (WhatmanTM, GE Healthcare, Freiburg, Germany). Each filter was washed with 5 ml of 25 mM Tris base pH 8.0,100 mM NaCl, 5 mM MgCl_2_, suspended in 5 ml of Ultima GoldTM (Perkin Elmer, Waltham, MA, USA) solution, and measured in a Tri-Carb 2810 TR scintillation analyzer (Perkin Elmer, Waltham, MA, USA). The assay was adapted to the matrix-bound TAP-Cdc24 by mixing the magnetic beads with assay buffer, premixed with Cdc42 and [αP32]GTP. For each time-point 25 µl of the mixture was removed and placed in a magnetic rack to precipitate the beads. The supernatant was subsequently quenched in 500 µl of ice-cold assay buffer and processed as described above.

### SPR measurements

6His-Cdc24_428-854_ and its mutants were cloned into a pET15b derived expression plasmid and expressed in *E.coli* BL21DE3 in super broth medium at 18°C. The recombinant proteins were purified from the crude extract by IMAC and size exclusion chromatography and transferred into HBSEP buffer (15mM HEPES, 150 mM NaCl, 3 mM EDTA, 0.05% Tween, pH 7.4) (Renz et al., 2013).

For comparing the binding of 6His-Cdc24_428-854_ to SNAP-tagged septin rods and SNAP-tagged Cdc11, the SNAP-tagged proteins were biotinylated with BG-Biotin, and subsequently immobilized on a CM5 chip (GE Healthcare, Freiburg, Germany) that displayed a covalently linked anti-Biotin antibody (US Biologicals, Salem, MA, USA) (Renz et al., 2013; Gronemeyer et al., 2016). Binding of purified 6His-Cdc24_428-854_ was recorded on a Biacore X-100, and the equilibrium binding constants (K_D_) were subsequently determined by the X100 evaluation software (GE Healthcare, Freiburg, Germany). All measurements were performed at least as triplicate.

For comparing the binding of 6His-Cdc24_428-854_, Cdc24_428-854K801A_, Cdc24_428-854 K525E_, and Cdc24_428-854 K525E K801A_ to Cdc11, 6His-Cdc11-SNAP was biotinylated with BG-Biotin and subsequently immobilized on a CAP chip (Biotin Capture Kit; GE Healthcare, Freiburg, Germany). Preparation of the chip, capture of the ligand, and regeneration of the chip surface was carried out according to the manufacturer’s recommendations. Purified 6His-Cdc24_428-854_ and its mutants were adjusted to the same concentrations in HBSEP buffer, and their binding to immobilized 6His-Cdc11-SNAP was recorded on the Biacore X-100. Measurements were performed as triplicate with comparable outcomes.

### GST-Cdc11 pull-down

*CDC11* was cloned in frame with GST into the pGEX2T plasmid. The protein was expressed in LB medium at 37 °C after addition of 0.1 mM IPTG. GST-Cdc11 and GST alone were immobilized on 100 µl equilibrated Glutathione sepharose slurry (GE Healthcare, Freiburg, Germany) directly from the crude extract. 1 µM of 6His-Cdc24_428-854_, or 6His-Bem1_PB_-SNAP in PBS were added, and the beads were washed and bound protein was eluted with an excess of Glutathione after 1 h incubation at 8 °C under gentle agitation. The eluates were separated by SDS-PAGE, blotted onto nitrocellulose and the proteins were detected with an anti-His antibody (Sigma-Aldrich, Steinheim, Germany). To test whether 6His-PB_Bem1_ and Cdc11 can bind simultaneously to Cdc24_428-854_, 2 µM or 4 µM of purified 6His-PB_Bem1_-SNAP were added to 6His-Cdc24_428-854_ pre-incubated GST-Cdc11-coated beads.

### Fluorescence microscopy

For microscopic inspection yeast cells were grown overnight in SD media, diluted 1:8 in 3-4 ml fresh SD medium, and grown for 3 to 6 h at 30 °C to mid-log phase. About 1 ml culture was spun down, and the cell pellet resuspended in 20-50 µl residual medium. 3 µl were spotted onto a microscope slide, the cells were immobilized with a coverslip and inspected under the microscope. For time-resolved imaging 3 µl of prepared cell suspension was mounted on a SD-agarose pad (1.7% agarose), embedded in a customized glass slide, and sealed by a cover slip fixed by parafilm stripes. Imaging was started after 15 to 30 min recovery at 30 °C. SPLIFF and other time-lapse experiments were observed with a DeltaVision wide-field fluorescence microscope system (GE Healthcare, Freiburg, Germany) provided with a Olympus IX71 microscope, a steady-state heating chamber, a CoolSNAP HQ2 and CascadeII512-CCD camera both by Photometrics (Tucson, AZ, USA), a U Plan S Apochromat 100Å∼ 1.4 NA oil **∞**/0.17/FN26.5 objective and a Photofluor LM-75 halogen lamp (Burlington, VT, USA). Images were visualized using softWoRx software (GE Healthcare) and adapted z series at 30°C. Exposure time was adapted tot he intensity of GFP and mCherry signal for every fluorescently labeled protein to reduce bleaching and phototoxicity. For further analyses an Axio Observer spinning-disc confocal microscope (Zeiss, Göttingen, Germany), equipped with an Evolve512 EMCCD camera (Photometrics, Tucson, AZ, USA), a Plan-Apochromat 63Å∼/1.4 oil DIC objective, and 488 nm and 561 nm diode lasers (Zeiss, Göttingen, Germany) was used. Images were analyzed with the ZEN2 software (Zeiss, Göttingen, Germany).

### Quantitative analysis of microscopy data and SPLIFF measurements

Microscopy data were processed and analyzed using ImageJ64 1.49 software and Excel. Analysis of temporal and spatial characteristics of the Cdc11-Cdc24 interaction by SPLIFF was performed as described (Moreno et al., 2013; Dunkler et al., 2015). *CDC11CCG* under its native promoter and *P_CUP1_-N_ub_-CDC24* were co-expressed in a-cells and grown in SD medium without copper sulfate. Cells were immobilized on an agarose pad and interactions were monitored by three channel z-stacks (5×0.6 µm) microscopy every 2 min. Z-slices with Cdc11 signals were projected by SUM-projection. The fluorescence intensities (FI) of mCherry and GFP channels were determined by integrated density measurements of the region of interest and a region within the cytosol. For each time point and channel the intracellular background was subtracted from the localized signal to obtain the localized fluorescence intensity (FI_red_ and FI_green_). The values were normalized to the mother bud neck signal after cytokinesis. For steady state SPLIFF measurements during bud emergence GFP- and mCherry-intensities were normalized to the Cdc11CCG bud neck signal of daughter cells. The resulting relative fluorescence intensity RFI(t) was then used to calculate the conversion FD(t):

FD(t)=RFI_red_-RFI_green_/RFI_red_

FD(t) as a readout of CCG- to CC conversion describes its temporal progress in percent. Excel was utilized for initial calculations, Prism 7.0 software (GraphPad) to plot the final graph. All error bars indicate SEM.

### Statistical evaluation

GraphPad Prism7 was applied for statistical data evaluation. Students *t*-tests were used to compare the percentage of cells of a certain phenotype. If the data sets did not follow a normal distribution the significance of differences between data were evaluated by Mann-Whitney-U-tests.

### Electron microscopy

Saturated cultures of the cells were diluted in fresh media and grown at 30 °C until an OD_600_ of 0.9-1.0. Cells were washed in PBS, attached to Poly-Lysine treated silicon wavers, and fixed with 2.5% glutaraldehyde (1% saccharose in PBS) for 1h at RT. Cells were post-fixed with Os_4_O_4_ (2% in PBS) and after washing in PBS gradually dehydrated in 30%, 50%, 790%, 90% and 100% propanol. Fixed cells were critically point dried using carbon dioxide. The samples were finally rotary-coated in a BAF 300 freeze etching device (Bal-tec, Liechtenstein) by electron beam evaporation with 3 nm of platinium carbon from an angle of 45°. Samples were inspected with a Hitachi S-5200 in-lens field emission SEM at an acceleration voltage of 2 kV (Hitachi, Tokio, Japan).

## Acknowledgements

We thank Steffi Timmermann and Ute Nussbaumer for technical assistance. We thank Judith Müller for advice during Cdc24 purification. The work was funded by grants from the DFG to N.J. (Jo 187/5-2). The authors declare no conflicts of interest.

## Author contribution

J. Chollet, A. Dünkler, T. Gronemeyer, N. Johnsson designed and analyzed the experiments. J. Chollet, and A. Dünkler performed the microscopic analysis of wild type- and *cdc24kk* cells. A. Dünkler performed the SPLIFF analysis. J. Chollet perfomed the EM analysis. J. Chollet and T. Gronemeyer performed the biochemical characterizations. L. Vivero-Pol performed the initial Split-Ub interaction screen. A. Bäuerle mapped the binding sites of Cdc11 on Cdc24. N. Johnsson conducted the genetic interaction analysis. N. Johnsson, A. Dünkler and J. Chollet designed the study. N. Johnsson together with the help of A. Dünkler, T. Gronemeyer and J. Chollet wrote the manuscript.

## References

Atkins, B.D., S. Yoshida, K. Saito, C.F. Wu, D.J. Lew, and D. Pellman. 2013. Inhibition of Cdc42 during mitotic exit is required for cytokinesis. J Cell Biol. 202:231–40.

Bähler, J., J.Q. Wu, M.S. Longtine, N.G. Shah, A. McKenzie 3rd, A.B. Steever, A. Wach, P. Philippsen, and J.R. Pringle. 1998. Heterologous modules for efficient and versatile PCR-based gene targeting in Schizosaccharomyces pombe. Yeast. 14:943–951.

Barral, Y., V. Mermall, M.S. Mooseker, and M. Snyder. 2000. Compartmentalization of the cell cortex by septins is required for maintenance of cell polarity in yeast. Mol.Cell. 5:841–851.

Bender, A., and J.R. Pringle. 1989. Multicopy suppression of the cdc24 budding defect in yeast by CDC42 and three newly identified genes including the ras-related gene RSR1. Proc Natl Acad Sci U S A. 86:9976–80.

Bertin, A., M.A. McMurray, P. Grob, S.S. Park, G. Garcia 3rd, I. Patanwala, H.L. Ng, T. Alber, J. Thorner, and E. Nogales. 2008. Saccharomyces cerevisiae septins: supramolecular organization of heterooligomers and the mechanism of filament assembly. Proc Natl Acad Sci U S A. 105:8274–9.

Bose, I., J.E. Irazoqui, J.J. Moskow, E.S. Bardes, T.R. Zyla, and D.J. Lew. 2001. Assembly of scaffold-mediated complexes containing Cdc42p, the exchange factor Cdc24p, and the effector Cla4p required for cell cycle-regulated phosphorylation of Cdc24p. J Biol Chem. 276:7176–86.

Brausemann, A., S. Gerhardt, A.K. Schott, O. Einsle, A. Grosse-Berkenbusch, N. Johnsson, and T. Gronemeyer. 2016. Crystal structure of Cdc11, a septin subunit from Saccharomyces cerevisiae. J.Struct.Biol. 193:157–161.

Brown, J.L., M. Jaquenoud, M.P. Gulli, J. Chant, and M. Peter. 1997. Novel Cdc42-binding proteins Gic1 and Gic2 control cell polarity in yeast. Genes Dev. 11:2972–2982.

Butty, A.C., N. Perrinjaquet, A. Petit, M. Jaquenoud, J.E. Segall, K. Hofmann, C. Zwahlen, and M. Peter. 2002. A positive feedback loop stabilizes the guanine-nucleotide exchange factor Cdc24 at sites of polarization. Embo J. 21:1565–1576.

Caviston, J.P., M. Longtine, J.R. Pringle, and E. Bi. 2003. The role of Cdc42p GTPase-activating proteins in assembly of the septin ring in yeast. Mol Biol Cell. 14:4051–66.

Chant, J., and I. Herskowitz. 1991. Genetic control of bud site selection in yeast by a set of gene products that constitute a morphogenetic pathway. Cell. 65:1203–1212.

Chiou, J.G., M.K. Balasubramanian, and D.J. Lew. 2017. Cell Polarity in Yeast. Annu.Rev.Cell Dev.Biol. 33:77–101.

Dohmen, R.J., R. Stappen, J.P. McGrath, H. Forrova, J. Kolarov, A. Goffeau, and A. Varshavsky. 1995. An essential yeast gene encoding a homolog of ubiquitin-activating enzyme. J Biol Chem. 270:18099–109.

Dowell, R.D., O. Ryan, A. Jansen, D. Cheung, S. Agarwala, T. Danford, D.A. Bernstein, P.A. Rolfe, L.E. Heisler, B. Chin, C. Nislow, G. Giaever, P.C. Phillips, G.R. Fink, D.K. Gifford, and C. Boone. 2010. Genotype to phenotype: a complex problem. Science. 328:469.

Dünkler, A., J. Müller, and N. Johnsson. 2012. Detecting protein-protein interactions with the split-ubiquitin sensor. Methods Mol Biol. 786:115–30.

Dünkler, A., R. Rosler, H.A. Kestler, D. Moreno-Andres, and N. Johnsson. 2015. SPLIFF: A Single-Cell Method to Map Protein-Protein Interactions in Time and Space. Methods Mol.Biol. 1346:151–168.

Evangelista, M., K. Blundell, M.S. Longtine, C.J. Chow, N. Adames, J.R. Pringle, M. Peter, and C. Boone. 1997. Bni1p, a yeast formin linking cdc42p and the actin cytoskeleton during polarized morphogenesis. Science. 276:118–122.

Fairn, G.D., M. Hermansson, P. Somerharju, and S. Grinstein. 2011. Phosphatidylserine is polarized and required for proper Cdc42 localization and for development of cell polarity. Nat.Cell Biol. 13:1424–1430.

Garcia, G., 3rd, A. Bertin, Z. Li, Y. Song, M.A. McMurray, J. Thorner, and E. Nogales. 2011. Subunit-dependent modulation of septin assembly: budding yeast septin Shs1 promotes ring and gauze formation. J Cell Biol. 195:993–1004.

Gladfelter, A.S., I. Bose, T.R. Zyla, E.S. Bardes, and D.J. Lew. 2002. Septin ring assembly involves cycles of GTP loading and hydrolysis by Cdc42p. J Cell Biol. 156:315–26.

Goryachev, A.B., and A.V. Pokhilko. 2008. Dynamics of Cdc42 network embodies a Turing-type mechanism of yeast cell polarity. FEBS Lett. 582:1437–1443.

Gronemeyer, T., J. Chollet, S. Werner, O. Glomb, A. Bauerle, and N. Johnsson. 2016. A Split-Ubiquitin Based Strategy Selecting for Protein Complex-Interfering Mutations. G3 (Bethesda). 6:2809–2815.

Gulli, M.P., M. Jaquenoud, Y. Shimada, G. Niederhauser, P. Wiget, and M. Peter. 2000. Phosphorylation of the Cdc42 exchange factor Cdc24 by the PAK-like kinase Cla4 may regulate polarized growth in yeast. Mol.Cell. 6:1155–1167.

Howell, A.S., and D.J. Lew. 2012. Morphogenesis and the cell cycle. Genetics. 190:51–77.

Howell, A.S., N.S. Savage, S.A. Johnson, I. Bose, A.W. Wagner, T.R. Zyla, H.F. Nijhout, M.C. Reed, A.B. Goryachev, and D.J. Lew. 2009. Singularity in polarization: rewiring yeast cells to make two buds. Cell. 139:731–43.

Hruby, A., M. Zapatka, S. Heucke, L. Rieger, Y. Wu, U. Nussbaumer, S. Timmermann, A. Dünkler, and N. Johnsson. 2011. A constraint network of interactions: protein-protein interaction analysis of the yeast type II phosphatase Ptc1p and its adaptor protein Nbp2p. J Cell Sci. 124:35–46.

Irazoqui, J.E., A.S. Gladfelter, and D.J. Lew. 2003. Scaffold-mediated symmetry breaking by Cdc42p. Nat.Cell Biol. 5:1062–1070.

Ito, T., Y. Matsui, T. Ago, K. Ota, and H. Sumimoto. 2001. Novel modular domain PB1 recognizes PC motif to mediate functional protein-protein interactions. Embo J. 20:3938–3946.

Iwase, M., J. Luo, S. Nagaraj, M. Longtine, H.B. Kim, B.K. Haarer, C. Caruso, Z. Tong, J.R. Pringle, and E. Bi. 2006. Role of a Cdc42p effector pathway in recruitment of the yeast septins to the presumptive bud site. Mol Biol Cell. 17:1110–25.

Janke, C., M.M. Magiera, N. Rathfelder, C. Taxis, S. Reber, H. Maekawa, A. Moreno-Borchart, G. Doenges, E. Schwob, E. Schiebel, and M. Knop. 2004. A versatile toolbox for PCR-based tagging of yeast genes: new fluorescent proteins, more markers and promoter substitution cassettes. Yeast. 21:947–62.

Johnsson, N., and A. Varshavsky. 1994. Split ubiquitin as a sensor of protein interactions in vivo. Proc Natl Acad Sci U S A. 91:10340–4.

Kadota, J., T. Yamamoto, S. Yoshiuchi, E. Bi, and K. Tanaka. 2004. Septin ring assembly requires concerted action of polarisome components, a PAK kinase Cla4p, and the actin cytoskeleton in Saccharomyces cerevisiae. Mol Biol Cell. 15:5329–45.

Kang, P.J., A. Sanson, B. Lee, and H.O. Park. 2001. A GDP/GTP exchange factor involved in linking a spatial landmark to cell polarity. Science. 292:1376–1378.

Kozubowski, L., K. Saito, J.M. Johnson, A.S. Howell, T.R. Zyla, and D.J. Lew. 2008. Symmetry-breaking polarization driven by a Cdc42p GEF-PAK complex. Curr Biol. 18:1719–26.

Kuo, C.C., N.S. Savage, H. Chen, C.F. Wu, T.R. Zyla, and D.J. Lew. 2014. Inhibitory GEF phosphorylation provides negative feedback in the yeast polarity circuit. Curr.Biol. 24:753–759.

Kustermann, J., Y. Wu, L. Rieger, D. Dedden, T. Phan, P. Walther, A. Dunkler, and N. Johnsson. 2017. The cell polarity proteins Boi1p and Boi2p stimulate vesicle fusion at the plasma membrane of yeast cells. J.Cell.Sci. 130:2996–3008.

Laan, L., J.H. Koschwanez, and A.W. Murray. 2015. Evolutionary adaptation after crippling cell polarization follows reproducible trajectories. Elife. 4:10.7554/eLife.09638.

Lai, H., J.G. Chiou, A. Zhurikhina, T.R. Zyla, D. Tsygankov, and D.J. Lew. 2018. Temporal regulation of morphogenetic events in Saccharomyces cerevisiae. Mol.Biol.Cell. 29:2069–2083.

Lamson, R.E., M.J. Winters, and P.M. Pryciak. 2002. Cdc42 regulation of kinase activity and signaling by the yeast p21-activated kinase Ste20. Mol.Cell.Biol. 22:2939–2951.

Laser, H., C. Bongards, J. Schuller, S. Heck, N. Johnsson, and N. Lehming. 2000. A new screen for protein interactions reveals that the Saccharomyces cerevisiae high mobility group proteins Nhp6A/B are involved in the regulation of the GAL1 promoter. Proc Natl Acad Sci U S A. 97:13732–7.

Longtine, M.S., A. McKenzie 3rd, D.J. Demarini, N.G. Shah, A. Wach, A. Brachat, P. Philippsen, and J.R. Pringle. 1998. Additional modules for versatile and economical PCR-based gene deletion and modification in Saccharomyces cerevisiae. Yeast. 14:953–961.

Marquardt, J., X. Chen, and E. Bi. 2019. Architecture, remodeling, and functions of the septin cytoskeleton. Cytoskeleton (Hoboken). 76:7–14.

McCusker, D., C. Denison, S. Anderson, T.A. Egelhofer, J.R. Yates 3rd, S.P. Gygi, and D.R. Kellogg. 2007. Cdk1 coordinates cell-surface growth with the cell cycle. Nat Cell Biol. 9:506–15.

Meca, J., A. Massoni-Laporte, D. Martinez, E. Sartorel, A. Loquet, B. Habenstein, and D. McCusker. 2019. Avidity-driven polarity establishment via multivalent lipid-GTPase module interactions. Embo J. 38:10.15252/embj.201899652. Epub 2018 Dec 17.

Mionnet, C., S. Bogliolo, and R.A. Arkowitz. 2008. Oligomerization regulates the localization of Cdc24, the Cdc42 activator in Saccharomyces cerevisiae. J.Biol.Chem. 283:17515–17530.

Moran, K.D., H. Kang, A.V. Araujo, T.R. Zyla, K. Saito, D. Tsygankov, and D.J. Lew. 2019. Cell-cycle control of cell polarity in yeast. J.Cell Biol. 218:171–189.

Moreno, D., J. Neller, H.A. Kestler, J. Kraus, A. Dünkler, and N. Johnsson. 2013. A fluorescent reporter for mapping cellular protein-protein interactions in time and space. Mol Syst Biol. 9:647.

Nagata, K., and M. Inagaki. 2005. Cytoskeletal modification of Rho guanine nucleotide exchange factor activity: identification of a Rho guanine nucleotide exchange factor as a binding partner for Sept9b, a mammalian septin. Oncogene. 24:65–76.

Neller, J., A. Dunkler, R. Rosler, and N. Johnsson. 2015. A protein complex containing Epo1p anchors the cortical endoplasmic reticulum to the yeast bud tip. J.Cell Biol. 208:71–87.

Okada, S., M. Leda, J. Hanna, N.S. Savage, E. Bi, and A.B. Goryachev. 2013. Daughter cell identity emerges from the interplay of cdc42, septins, and exocytosis. Dev Cell. 26:148–61.

Orlando, K., J. Zhang, X. Zhang, P. Yue, T. Chiang, E. Bi, and W. Guo. 2008. Regulation of Gic2 localization and function by phosphatidylinositol 4,5-bisphosphate during the establishment of cell polarity in budding yeast. J Biol Chem. 283:14205–12.

Park, H.O., E. Bi, J.R. Pringle, and I. Herskowitz. 1997. Two active states of the Ras-related Bud1/Rsr1 protein bind to different effectors to determine yeast cell polarity. Proc Natl Acad Sci U S A. 94:4463–8.

Peterson, J., Y. Zheng, L. Bender, A. Myers, R. Cerione, and A. Bender. 1994. Interactions between the bud emergence proteins Bem1p and Bem2p and Rho-type GTPases in yeast. J.Cell Biol. 127:1395–1406.

Pruyne, D., A. Legesse-Miller, L. Gao, Y. Dong, and A. Bretscher. 2004. Mechanisms of polarized growth and organelle segregation in yeast. Annu Rev Cell Dev Biol. 20:559–91.

Rapali, P., R. Mitteau, C. Braun, A. Massoni-Laporte, C. Unlu, L. Bataille, F.S. Arramon, S.P. Gygi, and D. McCusker. 2017. Scaffold-mediated gating of Cdc42 signalling flux. Elife. 6:10.7554/eLife.25257.

Reichel, C., and N. Johnsson. 2005. The split-ubiquitin sensor: measuring interactions and conformational alterations of proteins in vivo. Methods Enzymol. 399:757–776.

Renz, C., N. Johnsson, and T. Gronemeyer. 2013. An efficient protocol for the purification and labeling of entire yeast septin rods from E.coli for quantitative in vitro experimentation. BMC Biotechnol. 13:60.

Sadian, Y., C. Gatsogiannis, C. Patasi, O. Hofnagel, R.S. Goody, M. Farkasovsky, and S. Raunser. 2013. The role of Cdc42 and Gic1 in the regulation of septin filament formation and dissociation. Elife. 2:e01085.

Schaub, Y., A. Dunkler, A. Walther, and J. Wendland. 2006. New pFA-cassettes for PCR-based gene manipulation in Candida albicans. J.Basic Microbiol. 46:416–429.

Schneider, C., J. Grois, C. Renz, T. Gronemeyer, and N. Johnsson. 2013. Septin rings act as a template for myosin higher-order structures and inhibit redundant polarity establishment. J Cell Sci. 126:3390–3400.

Shimada, Y., P. Wiget, M.P. Gulli, E. Bi, and M. Peter. 2004. The nucleotide exchange factor Cdc24p may be regulated by auto-inhibition. Embo J. 23:1051–62.

Sikorski, R.S., and P. Hieter. 1989. A system of shuttle vectors and yeast host strains designed for efficient manipulation of DNA in Saccharomyces cerevisiae. Genetics. 122:19–27.

Sloat, B.F., A. Adams, and J.R. Pringle. 1981. Roles of the CDC24 gene product in cellular morphogenesis during the Saccharomyces cerevisiae cell cycle. J.Cell Biol. 89:395–405.

Sloat, B.F., and J.R. Pringle. 1978. A mutant of yeast defective in cellular morphogenesis. Science. 200:1171–1173.

Smith, S.E., B. Rubinstein, I. Mendes Pinto, B.D. Slaughter, J.R. Unruh, and R. Li. 2013. Independence of symmetry breaking on Bem1-mediated autocatalytic activation of Cdc42. J.Cell Biol. 202:1091–1106.

Versele, M., and J. Thorner. 2004. Septin collar formation in budding yeast requires GTP binding and direct phosphorylation by the PAK, Cla4. J Cell Biol. 164:701–15.

Wai, S.C., S.A. Gerber, and R. Li. 2009. Multisite phosphorylation of the guanine nucleotide exchange factor Cdc24 during yeast cell polarization. PLoS One. 4:e6563.

Wedlich-Soldner, R., S.C. Wai, T. Schmidt, and R. Li. 2004. Robust cell polarity is a dynamic state established by coupling transport and GTPase signaling. J Cell Biol. 166:889–900.

Weiss, E.L., A.C. Bishop, K.M. Shokat, and D.G. Drubin. 2000. Chemical genetic analysis of the budding-yeast p21-activated kinase Cla4p. Nat.Cell Biol. 2:677–685.

Witte, K., D. Strickland, and M. Glotzer. 2017. Cell cycle entry triggers a switch between two modes of Cdc42 activation during yeast polarization. Elife. 6:10.7554/eLife.26722.

Wittke, S., N. Lewke, S. Muller, and N. Johnsson. 1999. Probing the molecular environment of membrane proteins in vivo. Mol Biol Cell. 10:2519–30.

Woods, B., C.C. Kuo, C.F. Wu, T.R. Zyla, and D.J. Lew. 2015. Polarity establishment requires localized activation of Cdc42. J.Cell Biol. 211:19–26.

Woods, B., H. Lai, C.F. Wu, T.R. Zyla, N.S. Savage, and D.J. Lew. 2016. Parallel Actin-Independent Recycling Pathways Polarize Cdc42 in Budding Yeast. Curr.Biol. 26:2114–2126.

Yoshinaga, S., M. Kohjima, K. Ogura, M. Yokochi, R. Takeya, T. Ito, H. Sumimoto, and F. Inagaki. 2003. The PB1 domain and the PC motif-containing region are structurally similar protein binding modules. Embo J. 22:4888–4897.

